# Stress-induced brain extracellular vesicles ameliorate anxiety behaviour

**DOI:** 10.1101/2024.05.16.594479

**Authors:** Yusuke Mizohata, Yusuke Yoshioka, Minori Koga, Hiroyuki Toda, Hiroyuki Ohta, Yasunobu Kobayashi, Takahiro Ochiya, Yuji Morimoto

**Affiliations:** Department of Physiology, National Defense Medical College, Tokorozawa, Saitama, Japan; Aeromedical Laboratory, Japan Air Self-Defense Force, Sayama, Saitama, Japan; Department of Molecular and Cellular Medicine, Institute of Medical Science, Tokyo Medical University, Shinjuku, Tokyo, Japan; Department of Psychiatry, National Defense Medical College, Tokorozawa, Saitama, Japan; Department of Pharmacology, National Defense Medical College, Tokorozawa, Saitama, Japan; NanoSomiX Japan, Inc., Shinjuku, Tokyo, Japan

## Abstract

Extracellular vesicles (EVs), nano particles secreted by all types of cells, serve as a communication network, carrying information through the bloodstream to distant cells^1,2^. Notably, brain cells secrete EVs that play a crucial role in regulating neurological functions^3-5^. Meanwhile the brain detects acute stress and activates mechanisms to enhance stress resistance and maintain homeostasis^6,7^. However, the specific contribution of brain-derived extracellular vesicles (BDEVs) in modulating the stress response remains elusive. Here we found that administration of the acute stress-induced BDEVs to mice reduced anxiety-related behaviours, and this reduction was also induced by the administration of only three microRNAs (miRNAs) (miR-199a-3p, miR-99b-3p and miR-140-5p) included in the acute stress-induced BDEVs. Furthermore, we showed that miR-199a-3p contributes to the mechanism of the anxiolytic effect through the suppression of *Mecp2* in neurons. These findings elucidate the role of BDEVs in modulating mental activity under acute stress and provide insights into the underlying molecular mechanisms. Our results open up new avenues for therapeutic strategies in the treatment of anxiety disorders using EVs or miRNAs.

## Introduction

Extracellular vesicles (EVs), typically around 100 nm in size, are secreted by all cell types in the body and are carried by body fluids. EVs play a crucial role in transmitting molecular information to distant cells. These vesicles encapsulate physiologically active molecules, including proteins, mRNAs and microRNAs (miRNAs), originating from their parent cells^1,2^. Among the different types of Evs found in body fluids, those secreted by brain cells (Brain-derived extracellular vesicles, BDEVs)^4^ have been shown to play important roles in maintaining brain function, such as synaptic transmission^5^ and delivery of neurotrophic factors^3^.

Meanwhile, the brain is the centre of the stress response, but little is known about the specific functions of BDEVs in this process. It is well known that the stress response of the body initially enhances physiological resistance in acute phase, followed by a decrease in this resistance during chronic phase^6,7^. Giving that EVs mirror the states of the cells they originated from^8^, BDEVs secreted during the acute phase might exhibit an anti-stress effect. However, the specific physiological effects of these BDEVs during acute stress phase remain unclear. On the other hand, miRNAs that are included in BDEVs, are known to be fine tuners in the regulation of gene expression^9,10^ and thus their potential significance in the stress response is noteworthy, yet their roles in the acute phase is currently unknown.

Therefore, we aimed to investigate the anti-stress effects of BDEVs secreted during the acute phase of stress and to identify the key miRNAs responsible for these effects.

## Results

### Evaluation of the isolated BDEVs and miRNAs

The BDEVs were isolated from serum samples through a process involving ultracentrifugation and immunoprecipitation using anti-CD171 antibody (**Fig. 1a**). Subsequently, the quality of the isolated BDEVs was evaluated.

**Fig. 1.**
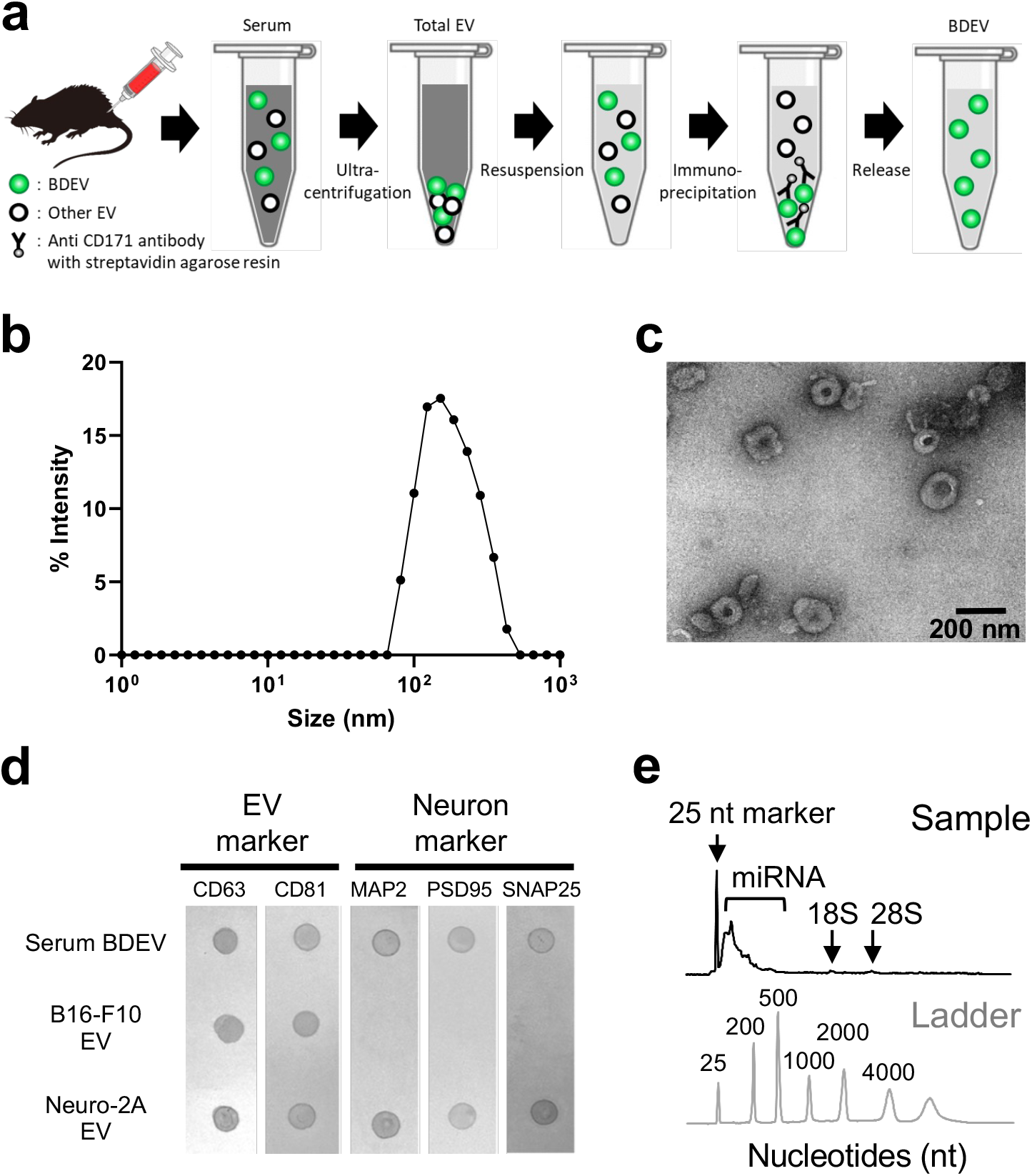
Evaluation of the isolated BDEVs and miRNAs. **a**, Overview of methods for isolation of BDEVs from serum. **b**,**c**, Analysis of total EV suspensions by dynamic light scattering (***b***) and transmission electron microscopy(***c***) revealed a particle size distribution of 50-200 nm and the presence of EV-like particles. **d**, Dot blotting analysis. BDEVs isolated from serum was equivalent to EVs isolated from neuronal (Neuro-2A) culture supernatant, but different from those obtained from non-neuronal cell (B16-F10, melanoma cell). Thus, BDEVs obtained from serum proved to be of brain origin. **e**, Electrophoresis analysis. The miRNAs extracted from BDEV were short chain and free of cell-derived rRNA contamination. EV, Extracellular vesicles. BDEV, Brain-derived extracellular vesicles.

First, the isolated particles were identified to EVs because they showed the EV-like particles with the size between 50-200 nm (**Fig. 1b, 1c**), presenting CD63-positive and CD81-positive (**Fig. 1d**). Further analysis by dot blotting confirmed the presence of neuronal markers in the isolated EVs (**Fig. 1d**) and showed that the luminescence intensity of these markers was dependent on EV concentration (**Extended Data Fig. 1**). These results indicate that the nano-size particles obtained by the approach described above were highly probable to be BDEVs.

The quality of the miRNAs extracted from BDEVs was also assessed, demonstrating a high level of purity, which was evidenced by the characteristic short-chain nucleotide composition and absence of cell-derived rRNAs (**Fig. 1e**).

### Stress alters BDEVs’ miRNA Profile

To demonstrate the response of brain cells to acute stress, the expression of *c-fos*, a classic marker of neuronal activity and immediate-early genes, was analysed. (**Fig. 2a**). The results showed that 1 h of water immersion restraint stress significantly increased *c-fos* mRNA expression in the hypothalamus (**Fig. 2b**). In contrast, acute stress did not alter the particle size distribution and morphological appearance (**Extended Data Fig. 2**). Similarly, the estimated levels of BDEV quantity, as determined by protein quantification, and the levels of miRNA concentration were not altered by acute stress, suggesting that this stress does not affect the BDEV secretion and the amount of miRNAs contained therein (**Fig. 2c,2d**).

**Fig. 2.**
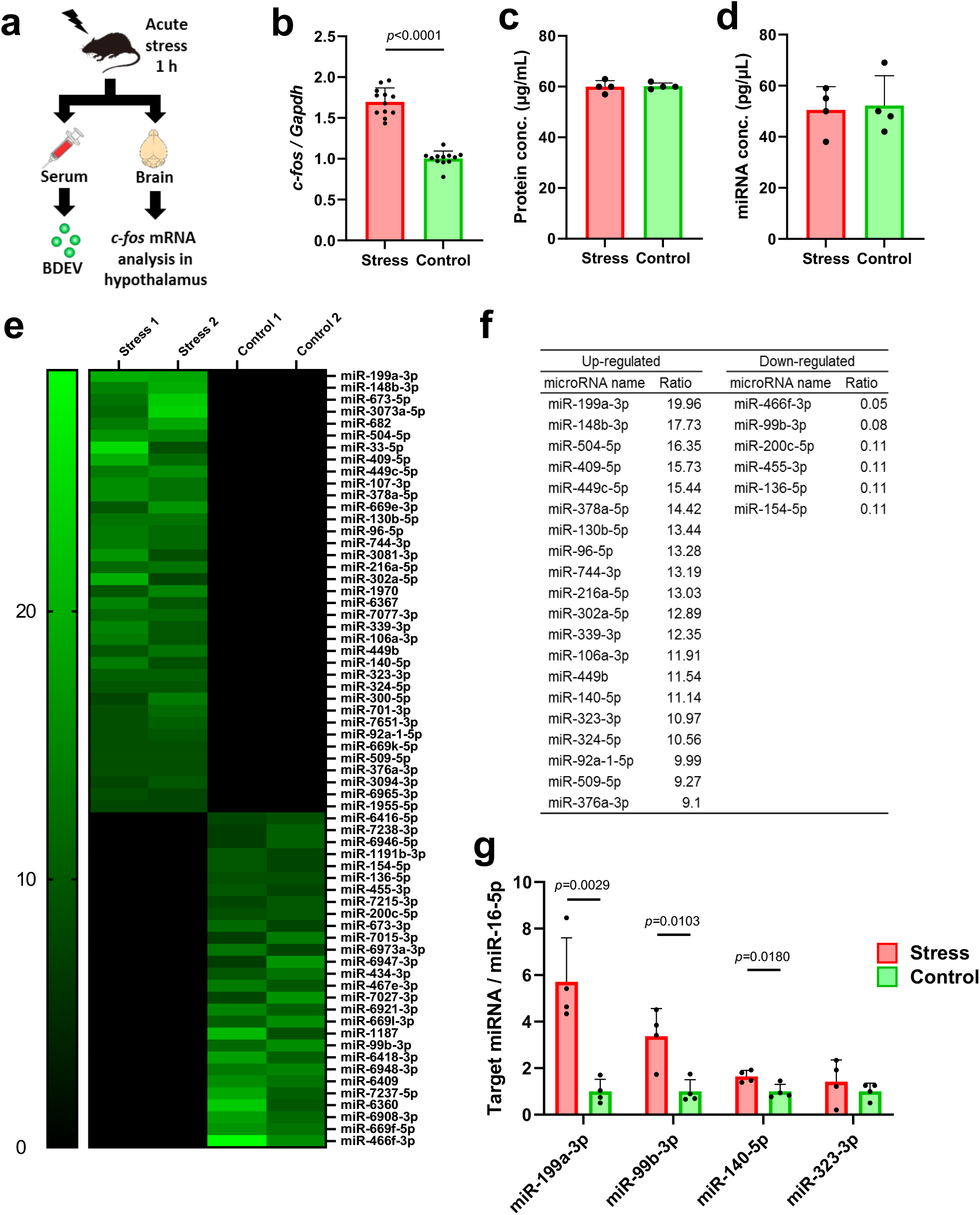
Acute stress alters the profile of miRNAs encapsulated in BDEVs. **a**, Protocol for tissue sampling after acute stress exposure. **b**, *c-fos* mRNA expression in the hypothalamus was increased by the acute stress (n=12). **c,d**, Protein concentrations in BDEVs suspensions (***c***) and concentration of miRNAs encapsulated in BDEVs (***d***) showed no change due to acute stress (n=4). Thus acute stress is considered to have no effect on BDEVs secretion. **e**, Microarrays analysis of BDEVs. 65 miRNAs in BDEVs were altered by acute stress (n=2 pools, each consisting of 8 mice). **f**, Extracted 26 miRNAs. Of the 65 miRNAs shown in ***e***, 26 miRNAs were identified as having human homologues. **g**, Extracted 4 miRNAs. Of the 26 miRNAs shown in ***f***, four miRNAs showed stable Ct values, three of which were significantly increased by acute stress (n=4 pools, each consisting of 2 mice). Data are mean±SD. Statistical analysis was performed using Student’s two-tailed *t*-tests. BDEV, Brain-derived extracellular vesicles.

Next, to elucidate the BDE V-encapsulated miRNAs that are altered by acute stress, we performed a comprehensive microarray analysis (1900 miRNAs). The analysis revealed that 65 miRNAs exhibited more than an 8-fold change in response to acute stress (**Fig. 2e and Supplementary Table 1**). Of these 65 miRNAs, 26 miRNAs with known human homologs (**Fig. 2f**) were validated by real-time RT-PCR. We found that four miRNAs (miR-199a-3p, miR-99b-3p, miR-140-5p and miR-323-3p) showed stable Ct values in both groups of samples (acute stress group and control group), and three of the miRNAs (miR-199a-3p, miR-99b-3p and miR-140-5p) demonstrated a significant elevation in 2^-ΔΔCt^ values in the acute stress group, while the fourth miRNA (miR-323-3p) showed no significant changes (**Fig. 2g**). Apart from these 4 miRNAs, the four other miRNAs (miR-376a-3p, miR-136-5p, miR-339-3p and miR-466f-3p) in the 26 miRNAs could not be accurately quantified (**Extended Data Fig. 3**). The remaining 18 miRNAs were not detected in both groups (**Supplementary Table 2**).

### Stress-induced BDEVs reduces anxiety-like behaviour

The effects of BDEVs and the miRNAs they contain on anxiety-related behaviour were assessed. Either BDEVs derived from stressed mice (stress-BDEVs), from control mice (control-BDEVs), or a vehicle solution was administered intravenously to a new group of mice that had been exposed to acute stress (**Fig. 3a, 3c**). In the open field test, the stress-BDEVs group showed a significant increase in time stayed in the centre while no significant differences were seen in the frequency of centre entry and in the total distance traveled among the three groups (**Fig. 3b (part of the Extended Data Fig. 4b extracted and reconstructed**)). Additionally, in the elevated plus-maze test, stress-BDEVs group showed a more frequent entry into the open arm and a greater distance travelled in the open arm while no significant differences were seen in the total distance travelled among the three groups (**Fig. 3d (part of the Extended Data Fig. 4c extracted and reconstructed**)). Given that mice exhibiting depressive-like symptoms typically show reduced activity in the central area of the open field test and the open arms of the elevated plus-maze^11^, the observed increase in activity in these areas suggests an anti-stress-like effect. This behavioural change suggests a potential reduction in depressive-like symptoms.

**Fig. 3.**
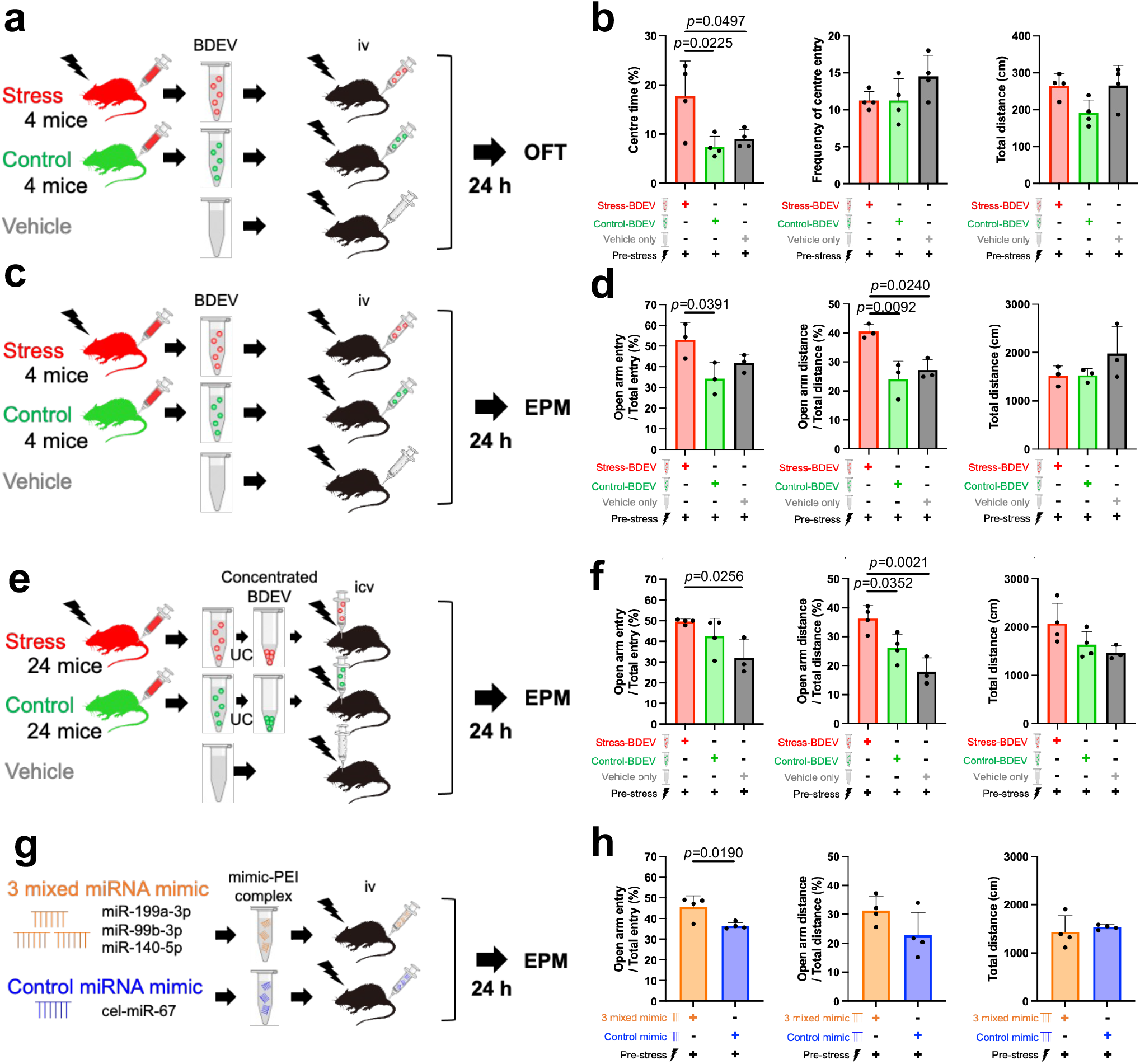
Stress-induced BDEVs and encapsulated miRNAs ameliorate anxiety behaviour. **a, c, e**, Experimental protocol and groups to assess the effects of BDEV administered intravenously (***a,c***) or intracerebroventricularly (***e***) on behavioural tests. **b**, Assessment by the open field test (OFT) regarding experiment ***a***. Intravenous administration of stress-BDEV increased the time spent in the centre (n=4). **d**, Assessment by the elevated plus maze test (EPM) regarding experiment ***c***. Intravenous administration of stress-BDEV increased the frequency of entry into the open arm and the distance travelled in the open arm (n=3). f, Assessment by the elevated plus maze test (EPM) regarding experiment ***e***. Intracerebroventricular administration of stress-BDEV increased the distance travelled in the open arm (n=3-4). **g**, Experimental protocol and groups to assess the effects of miRNA mimics administered intravenously on behavioural tests. **h**, Assessment by the elevated plus maze test (EPM) regarding experiment ***g***. Intravenous administration of 3 mixed miRNA mimic increased the frequency of entry into the open arm (n=4).

Since BDEVs showed anxiolytic effects, we hypothesised that their primary target might be the central nervous system. To test this hypothesis and determine whether BDEVs directly affect the brain, we injected them into the cerebral ventricles (**Fig. 3e**). Similar to the results observed with intravenous administration, in the elevated plus-maze test, mice in the stress-BDEVs group showed a significant increase in the distance travelled in the open arm when compared to both the control-BDEVs and vehicle groups (**Fig. 3f**). In contrast, there was no difference in total distance travelled among the groups (**Fig. 3f**).

Next, to determine whether the three stress-increased miRNAs (miR-199a-3p, miR-99b-3p and miR-140-5p) (**Fig. 2g)** induced behavioural changes, we injected a mixture of their mimics into mice intravenously and assessed behavioural effects (**Fig. 3g**). The mimic mixture group showed significantly more frequent entry into the open arm compared to the control group, but the total distance travelled was similar between groups (**Fig. 3h**).

### BDEVs’ miRNAs repress neuronal *Mecp2*

Since BDEVs was found to act on the central nervous system, we investigated their uptake into neural cells. Upon adding fluorescently labeled BDEVs to neuron (Neuro-2A cells), astrocyte (primary astrocytes), and microglia (BV-2 cells), fluorescent signals were obtained in each cell type, indicating that BDEVs was taken up by all cell types (**Fig. 4a**).

**Fig. 4.**
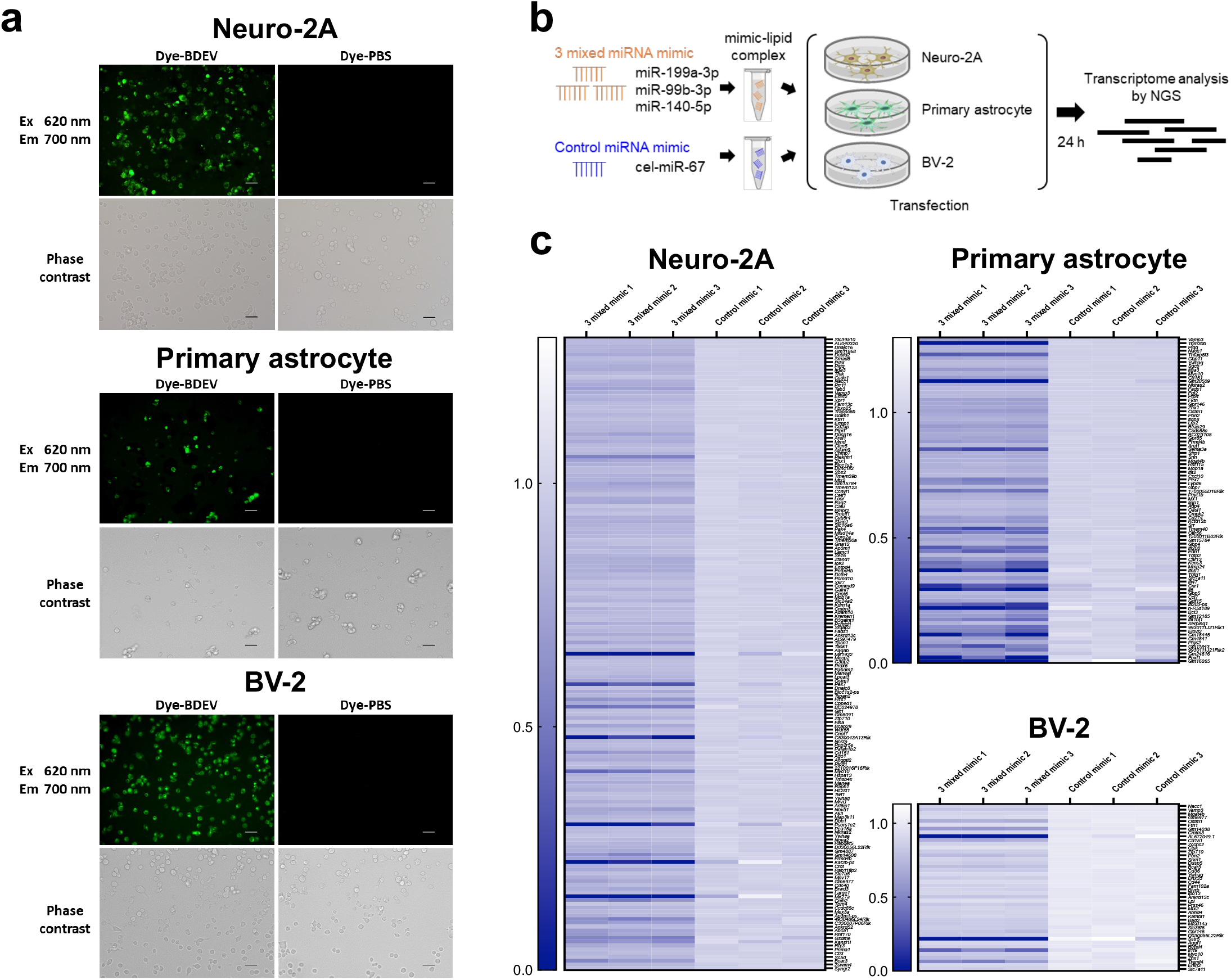
Stress-induced miRNAs alter mRNA profiles in neural cells. **a**, BDEV uptake by neural cells. BDEV fluorescently labelled with Mem Dye-Deep Red (EX03, Dojindo) from serum (8 mice) was uptaken by Neuro-2A, primary astrocyte and BV-2 (left-upper panel of each cell section). Conversely, no fluorescence signal was detected in all cells when the same procedure was performed in PBS with a fluorescent dye added (right-upper). The lower panel shows the phase contrast images. Scale bar=50 µm. **b**, Experimental protocol of transient transfection of stress-induced miRNA mimic into neural cells. **c**, Downregulated mRNAs revealed by NGS. Stress-induced 3 miRNAs (miR-199a-3p, miR-99b-3p and miR-140-5p) significantly decreased the expression of 167, 86 and 44 mRNAs in Neuro-2A, primary astrocyte and BV-2, respectively (n=3). Statistical analysis was performed using benjamini-hochberg method. BDEV, Brain-derived extracellular vesicles. NGS, Next generation sequencing.

Next, to identify target genes of stress-induced miRNAs in the neural cells, after transiently transfecting three mixed miRNA (miR-199a-3p, miR-99b-3p and miR-140-5p) mimics into neural cells (Neuro-2A, primary astrocytes and BV-2), we performed a comprehensive analysis of the repressed mRNAs using next-generation sequencing (NGS) (**Fig. 4b**). Consequently, the expression of 167, 86, and 44 mRNAs was found to be significantly downregulated in Neuro-2A, Primary astrocyte and BV-2, respectively (**Fig. 4c and Supplementary Table 3**). Among the down-regulated genes identified by NGS, target genes potentially regulated by the stress-induced miRNAs (miR-199a-3p, miR-99b-3p and miR-140-5p) were derived using TargetScanMouse databases (**Supplementary Table 4**-**6**), resulting in that 79 genes in Neuro-2A, 30 genes in primary astrocytes and 22 genes in BV-2 were identified (**Fig. 5a**). When these mRNAs, extracted from the Target Scan Mouse databases, were sorted according to the three miRNAs (miR-199a-3p, miR-99b-3p and miR-140-5p), the most dominant regulator was found to be miR-199a-3p in all three cell types, Neuro-2A, primary astrocytes and BV-2 (**Fig. 5b**).

**Fig. 5.**
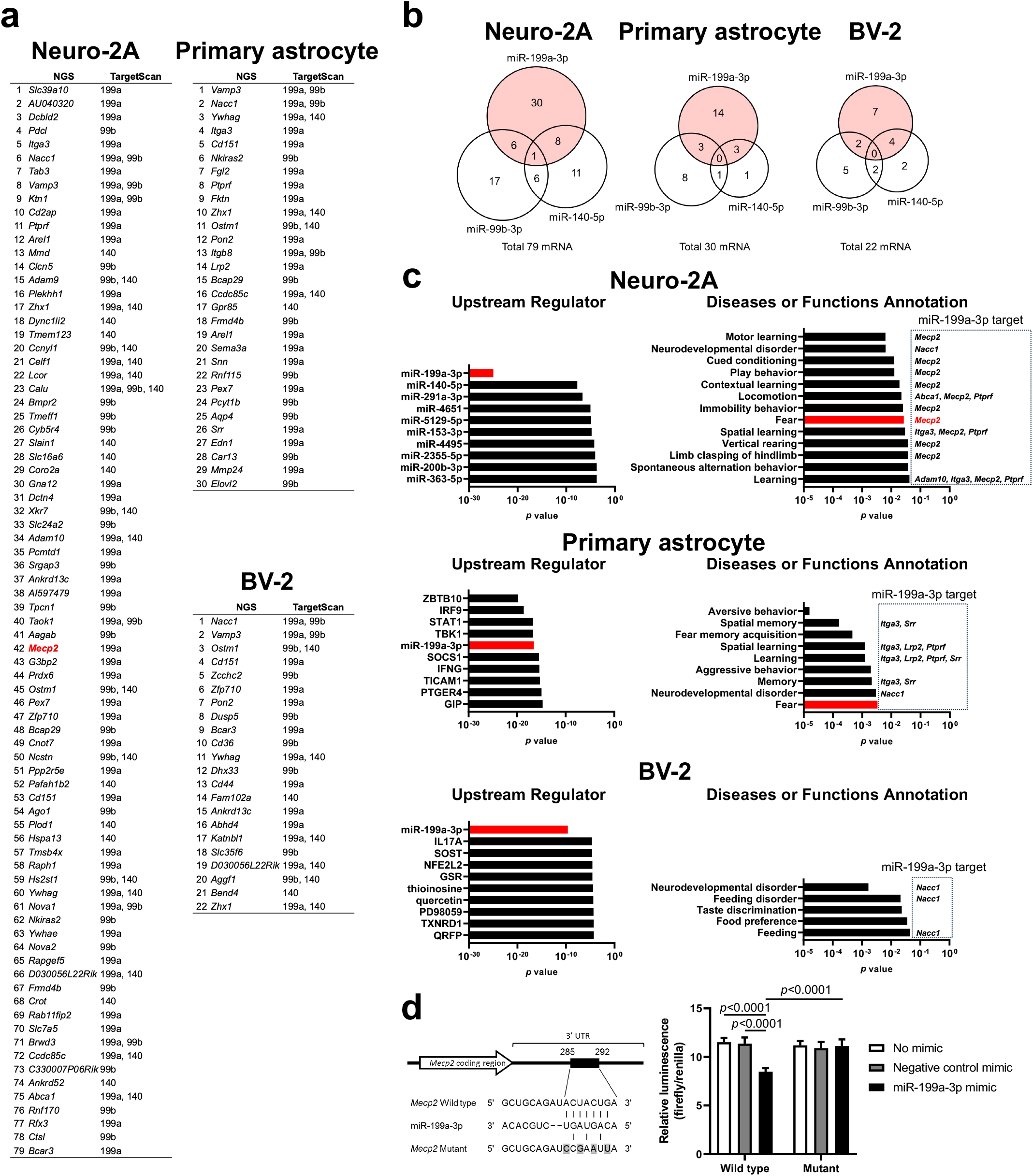
Stress-induced miRNA suppresses *Mecp2* expression in neuron. **a**, List of mRNAs directly targeted by miR-199a-3p, miR-99b-3p and/or miR-140-5p. Among the down-regulated mRNAs identified by NGS analysis (Fig. 4c), the number of mRNAs directly targeted by miR-199a-3p, miR-99b-3p and/or miR-140-5p was 79, 30 and 22 in Neuro-2A, primary astrocytes and BV-2, respectively. **b**, Categorisation of the mRNAs presented in result ***a*** broken down by target for each of the three miRNAs. Targets of miR-199a-3p was the most prevalent in all cell types. **c**, The left-hand graph shows the top 10 upstream regulators revealed by IPA applied for all down-regulated mRNAs. In all cell types, miR-199a-3p was the most dominant upstream regulator in the three miRNAs. The graph on the right shows categories of behaviour that significantly changed, as identified by the Disease or Function Annotation analysis of IPA. In addition, the mRNAs shown within the *dashed lines* are those associated with each category and identified as targets of miR-199a-3p by TargetScanMouse. As a result, *Mecp2* was extracted as a Fear-related factor directly regulated by miR-199a-3p in Neuro-2A. **d**, Schematic diagram of the 3′ UTR reporter assay (left) and identification of the miR-199a-3p binding site in *Mecp2* (right). miR-199a-3p suppresses *Mecp2* expression by binding to position 285-292 in its 3’UTR. Data are mean±SD. Statistical analysis was performed using one-way ANOVA with Tukey-Kramer post hoc test and Student’s two-tailed *t*-tests. NGS, Next generation sequencing. IPA, Ingenuity pathway analysis. UTR, Untranslated region.

Furthermore, by using Ingenuity Pathway Analysis (Qiagen) for Upstream Regulator analysis, we assessed the contribution of the three miRNAs (miR-199a-3p, miR-99b-3p and/or miR-140-5p) on the down-regulated genes. The analysis revealed that miR-199a-3p is a highly reliable upstream regulator. In particular, miR-199a-3p showed the lowest *p*-values in neuro-2A and BV-2, providing strong evidence for a causal relationship (**Fig. 5c and Supplementary Table 7**). In addition, when we analysed the down-regulated genes using the Diseases or Functions Annotation analysis of Ingenuity Pathway Analysis (IPA), specifically focusing on ‘behaviour’ categories, we identified ‘Fear’ – a behaviour closely linked to anxiety – in both Neuro-2A and primary astrocyte cells (**Fig. 5c and Supplementary Table 8**). Next, using the TargetScanMouse database (**Supplementary Table 4**), we searched for candidate genes directly targeted by miR-199a-3p from the down-regulated genes in the ‘Fear’ category in Neuro-2A cells, and only *Mecp2* was found to fall in the category (**Fig. 5c**). However, in primary astrocytes, no direct target candidates for miR-199a-3p were found (**Fig. 5c**). In addition, the absence of miR-99b-3p and miR-140-5p targets extracted by Target Scan Mouse (**Supplementary Table 5**, **6**) within the ‘Fear’ category of these neural cells was also confirmed (**Extended Data Fig. 5 and Supplementary Table 8**).

Based on these results, an experiment (3’UTR reporter assay) was conducted to verify that miR-199a-3p binds to *Mecp2* and represses its expression (**Fig. 5d**). The miR-199a-3p mimic transfected cells with a vector containing the wild-type 3’UTR sequence of *Mecp2* showed significantly reduced luminescence levels compared to the negative control mimic group (**Fig. 5d**). However, no reduction in luminescence levels by the miR-199a-3p mimic was observed in cells transfected with vectors containing the mutant 3’UTR sequence (**Fig. 5d**). These results indicate that miR-199a-3p directly binds to *Mecp2* and represses its expression.

## Discussion

In this study, we showed that acute stress increases the levels of three miRNAs (miR-199a-3p, miR-99b-3p and miR-140-5p) that are encapsulated in BDEVs. Furthermore, our findings indicate that BDEVs secreted under acute stress induce anti-anxiety behaviour and those three miRNAs play a key role in this effect. In addition, we found that suppression of *Mecp2* expression by miR-199a-3p in neurons is involved in the induction of anxiolytic behaviour.

Previous studies have shown that EVs may influence anxiety-related behaviours, but the EVs in many reports were obtained from chronically stressed or long-term depressed patients, and they are total EVs in the circulating blood, and have further been shown to act as an anxiety-exacerbating factor^12,13^. Our study is notable for concentrating on brain-derived EVs specifically collected during the acute phase of stress. EV contents mirror the molecular signature of the cells from origin^8^, and the brain, as the primary stress-sensing organ^6,7^, is thought to secrete EVs that reflect the stress state^8^. It is therefore suggested that during the acute stress response phase, BDEVs rich in stress resistance factors are secreted.

Our acute phase hypothesis is also supported by Wang, *et al*^14^, showing that EVs released from acutely depressed mice model by inflammatory mediation have antidepressant effects. However, their study differs from ours in that they used whole blood-derived EVs, and also identified a protein (Sig-1R) rather than a miRNA as the molecule of action. Since Sig-1R protein was not detected in our proteomic results of stress-BDEVs (**Extended Data Fig. 6 and Supplementary Table 9**), the Sig-1R alteration is likely to occur in EVs secreted from non-brain tissues.

The origin of BDEV secretion and the cells targeted need to be discussed to elucidate the mechanism of anxiolytic action of BDE V. We speculate that the secretory origins of the BDEVs in this study are mainly neuronal, also with some contribution from oligodendrocytes. This speculation arises from the fact that CD171, used for immunoprecipitation, is expressed in neurons and oligodendrocytes but not in astrocyte^15^. Additionally, since microglia, which are of mesodermal origin, exhibit a different expression pattern compared to other neural cells^16^, the CD171 expression level in microglia is considered to be low.

Conversely, the BDEVs were uptaken by Neuro-2A, astrocytes and BV-2, suggesting that these cell types are part of the target cells. The analysis using Target Scan Mouse and Upstream Regulator revealed that miR-199a-3p targeted the most candidate genes in all three cell types examined, and miR-199a-3p was identified as the most reliable upstream regulator. This suggest that miR-199a-3p plays a dominant role in modulating anxiolytic behaviour.

Furthermore, *Mecp2* in Neuro-2A was identified as a target of miR-199a-3p involved in anxiety and fear behaviour, with its binding site was pinpointed as 285-292 in its 3’UTR. Moreover, another report has also confirmed *Mecp2* as a direct target of miR-199a-3p^17^, supporting our findings. Therefore, it is considered that miR-199a-3p was transported by BDEVs to the neuron, where it induced anxiolytic behaviour via downregulation of *Mecp2* expression. Indeed, our results are consistent with observations of heightened anxious behaviour in transgenic rats overexpressing of *Mecp2*^18^ and increased in anxiolytic behaviour due to mutations in *Mecp2*^19^.

Additional analyses of the target pathways for the three miRNAs using Target Scan Mouse (**Supplementary Table 4**-**6**) and Reactome Pathway Database (**Supplementary Table 10-12**) revealed that miR-199a-3p acts in a distinct pathway from the other two miRNAs (**Extended Data Fig. 7 a-c**). Specifically, miR-199a-3p is probably a direct target of factors on the PI3K/Akt/mTOR pathway, a critical cascade in the regulation of various cellular processes (controlling cell growth, cell survival, protein synthesis, and metabolism) (**Extended Data Fig. 7a**). This was further confirmed by qPCR, revealing that miR-199a-3p suppresses factors related to the PI3K/Akt/mTOR pathway in the neurons (**Extended Data Fig. 8a and Supplementary Table 13**). Consequently, these evidences suggest that miR-199a-3p plays a critical role in inhibition of the mTOR pathway (**Extended Data Fig. 8b)**. Previous studies in autism^20,21^, which often presents as a symptom of hyper-anxious behaviour show activation of the mTOR pathway and also amelioration of hyper-anxious behaviour by mTOR inhibitors^22^. In fact, mTOR has also been reported to be a direct target of miR-199a-3p^23^. Thus, inhibition of mTOR by miR-199a-3p may also contribute to the enhancement of anti-anxiety behaviour observed in this study.

This study suggested that astrocytes and microglia may contribute to anxiolytic behaviour, although they are not direct targets of miR-199a-3p.

For astrocytes, no direct target genes for miR-199a-3p were identified, even though they were classified in the “fear” and “fear memory acquisition” categories (**Fig. 5c**). This suggests that astrocytes may act via different mechanisms, such as regulation of neurotransmitter uptake^24,25^. On the other hand, microglia were not found to have a “fear” category (**Fig. 5c**), suggesting that they are less directly involved in anxiolytic behaviour, but may be involved in the clearance of extracellular EVs through nonselective macropinocytosis^26-28^. Neurotransmitter uptake by astrocytes^29,30^ and micropinocytosis in microglia^31^ are known to be promoted by enhancement of the PI3K/Akt/mTOR pathway, whereas this study demonstrated that miR-199a-3p suppresses PI3K/Akt/mTOR pathway-related factor in these cells by qPCR (**Extended Data Fig. 8a and Supplementary Table 13**). Thus, miR-199a-3p may contribute to anxiolytic behaviour by regulating the amount of extracellular neurotransmitters and EVs via suppression of the PI3K/Akt/mTOR pathway in glial cells and microglia (**Extended Data Fig. 8b**).

Based on these findings, we propose the following hypotheses (**Extended Data Fig. 9**): 1) acute stress increases miR-199a-3p levels in BDEVs; 2) these BDEVs are uptaken into neurons; 3) within these neurons, miR-199a-3p inhibits *Mecp2* ; 4) Anxiety behaviour is suppressed.

As a future perspective, further research will improve our understanding of the role and dynamics of BDEVs in stress responses, and may also pave the way for drug discovery utilizing BDEVs or miRNAs.

## Acknowledgements

We thank the following researchers for their support; M Fujita, D Saito and S Nakamura at the National Defense Medical College for facilities and equipment. T Kobayashi and S Hatakeyama at Saitama University for equipment support. M Ohgidani of Asahikawa Medical University provided cells. This study was supported by the following grants in Ministry of Defense; Advanced Defense Medical Research (Stress and Resilience) (FY 2016-FY 2022).

## Author contributions

Y. Mizohata and Y. Morimoto designed and conceived the experiments. Y. Y. performed RNA-Seq analysis and reporter assay. M.K. and H.T. supervised for behavioural test. H.O. performed intracerebroventricular administration. Y.K. performed dot blot analysis. T.O. supervised for experiments of extracellular vesicle and miRNA. Y.Mizohata wrote the manuscript with input from other authors supervised the study.

## Competing interests

The authors declare no competing interests.

## Methods

### Animals

The 8-week-old male C57BL/6J mice were purchased from CLEA Japan (Tokyo, Japan). Breeding conditions were 23 ± 1°C, 55 ± 10% relative humidity, light period 5:00-19:00, free feeding and drinking. All mice were used after acclimatisation to the breeding environment for one week. This study was approved by the National Defense Medical College Animal Ethics Review Committee (#19057).

### Brain-derived extracellular vesicles (BDEVs) isolation from Blood

We performed a two-step isolation of BDEVs from serum.

Initially, total extracellular vesicles (EVs) in serum were purified by ultracentrifugation. 0.8 mL of pooled serum from two mice was centrifuged (10,000 g, 10 min, 4 °C) to remove cell debris. The supernatant was transferred to ultracentrifuge tubes (344090, Beckman Coulter, Brea, CA, United States). Total EVs were precipitated using an Optima MAX ultracentrifuge (Beckman Coulter, Brea, CA, United States) with an MLS-50 swinging rotor at 110,000 g for 70 min at 4 °C. The acceleration and deceleration rates of the ultracentrifuge were set to the maximum settings of the equipment. A description of these detailed ultracentrifuge conditions is recommended in MISEV 2023^32^. Whole EVs pellet was resuspended in 250 µL of PBS containing protease/phosphatase inhibitor (78440, Thermo, Waltham, MA, United States).

Isolation of BDEVs was then performed following the method described in a previous study^33^ with modifications (**Fig. 1a**). Total EVs suspension was added 100 µL of 3 % BSA/PBS with 1.6 µg of biotinylated anti-human CD171 antibody (5 µg for microarray) (13-1719, eBioscience, San Diego, CA, United States), followed by shaking (1 h, 4 °C). Subsequently, 50 µL of 3 % BSA/PBS plus 40 µL of streptavidin agarose resin (125 µL for microarray) (20347, Thermo) was added and shaken (30 min, 4 °C). PBS 500 µL was added, and after centrifugation (200 g, 10 min, 4 °C), the supernatant was removed. The pellet was mixed with 50 µL of 0.05 M glycine-HCl (pH 2.8) for 10 s by a mixer. Then 5 µL of 1 M Tris-HCl (pH 8.0) was added and neutralised by mixing for a few seconds. After centrifugation (200 g, 5 min, 4 °C), the supernatant was collected. To estimate the BDEV concentration, a BCA Protein Assay (23225, Thermo) was performed.

### BDEVs identification

After the purification of total EVs, particle size distribution by dynamic light scattering (DLS) and morphological observations by transmission electron microscopy (TEM) were carried out. The DLS measurements (ELSZ-1000, Otsuka Electronics, Hirakata, Japan) were performed after agitation with a mixer to disperse EV aggregation. For TEM observations, the whole EVs suspension stained with 3% uranium acetate was placed on formvar-coated copper grids, and the specimens were then air-dried and examined. A JEM-1400plus (JEOL, Akishima, Japan) was set at 100 k V.

After the isolation of BDEVs by immunoprecipitation, EV and neuronal markers were detected by dot blotting. EVs purified from the culture supernatant of mouse neuroblastoma cells (Neuro-2A) (RCB2639, RIKEN Bioresource Research Centre, Tsukuba, Japan) were used as a positive control for brain cell-derived EVs, and EVs collected from the culture supernatant of mouse melanoma cells (B16-F10) (CRL-6322, ATCC, Gaithersburg, MD, United States) were used as a negative control. To estimate the amount of EVs in each sample, a mouse CD63-sandwich ELISA system was constructed and used to measure the relative amount of CD63. The concentration of EV suspension purified from Neuro-2A culture supernatant was defined as 100 U/mL and used as a quantification standard for the ELISA system. Based on this standard, the concentration of purified EV from each sample was adjusted to 1 U/mL. Each adjusted sample was treated with M-PER Mammalian Protein Extraction Reagent (78501, Thermo), after which a 3-fold serial dilution series starting from 1.0 mU was prepared and blotted onto nitrocellulose membranes. The membranes were then incubated with primary antibodies against neuron and EV markers (neuronal markers; MAP2 (GTX133109, GeneTex, Irvine, CA, United States), PSD95 (GTX133091, GeneTex), SNAP25 (GTX 113839, GeneTex), and EV markers; CD63 (#143902, Biolegend, San Diego, CA, United States), CD81 (#104901, Biolegend)). The membranes were then treated with secondary antibodies (biotinylated goat anti-rabbit IgG (#65-6140, Invitrogen, Rockford, IL, United States), biotinylated goat anti-rat IgG (#405428, Biolegend), biotinylated goat anti-hamster IgG (# 405501, Biolegend)), followed by incubation with HRP-conjugated streptavidin (Thermo) and TMB substrate (Moss, Pasadena, MA, United States). Characterization of EVs by these methods is encouraged by MISEV 2023^32^.

### Stress loading methods

Restraint water immersion stress was used for acute stress exposure to mice. Mice were confined in 25 mL tubes and submerged chest-deep in a 37°C water bath for 1 h (7:00-8:00). Control mice were transferred to a newly prepared cage where they were allowed free movement for 1 h during the same period.

### Sampling of specimens

After acute stress exposure, cardiac blood samples were collected under anesthesia (medetomidine hydrochloride/midazolam/butorphanol: 4/0.75/5 mg/kg body weight, administered intraperitoneally). The brain was then extracted after decapitation. Dissection was carried out rostrally at the optic chiasma and caudally at the border with the midbrain, and the block containing the hypothalamus was immersed in RNAlater Stabilization Solution (AM7020, Invitrogen, Waltham, MA, United States) (**Fig. 2a**). The immersed brain blocks were sectioned at the border of the right and left ridges of the hypothalamic papillae^34^ after overnighting at 4°C, and then stored at-80°C until analysis.

Blood was overnighted at 4°C, followed by serum separation through centrifugation at 4°C. The obtained serum was stored at-80°C until analysis.

### RNA extraction

RNA extraction from hypothalamus and cultured cells was performed using the miRNeasy Mini Kit (217004, Qiagen, Venlo, Netherlands) according to the instructions. Tissues or cells were homogenised with QIAzol lysis reagent (217004, Qiagen). Subsequently, chloroform was added and mixed. After centrifugation (12,000 g, 15 min, 4 °C), the upper layer was separated and mixed with 1.5 × volume of 100% ethanol. The sample solution was flowed through a MinElute spin column, allowing RNA adsorption, followed by washing with buffer, and then drying the column. Finally, RNase free water was added to the column. After centrifugation (16,100 g, 5 min), total RNA solution was obtained. The purity and concentration of the extracted total RNA solution were measured using Nanodrop (ND-1000, Thermo).

Extraction of BDEV-derived total RNA was performed using the miRNeasy Serum/Plasma Kit (217184, Qiagen) according to the instructions. The extraction procedure was similar to the method described above, with exceptions of performing the final column wash using 80% ethanol and quality-checking of miRNAs through a bioanalyzer (2100, Agilent Technologies, Santa Clara, CA, United States) using RNA6000 Pico Kit (5067-1513, Agilent Technologies).

### Microarray

RNA samples extracted from 2.5 mL pooled serum (eight mice) were quality-checked by a bioanalyzer, and submitted to a comprehensive analysis of 1900 miRNAs using the Mouse miRNA Oligo chip (3D-Gene, TORAY, TOKYO, Japan). The obtained results were analysed by Gene Spring GX 14.9.1.

### Real time RT-PCR

Reverse transcription and PCR were executed on total RNA extracted from the hypothalamus or cultured cells according to the following procedure. For the reverse transcription reaction, 500 ng of total RNA was used, and the reaction was conducted using SuperScript IV VILO Master Mix (11756050, Invitrogen). The reaction conditions were 16 °C for 30 min, 42 °C for 30 min and 85 °C for 5 min. PCR reactions were performed using TaqMan Universal Master Mix II (4440040, Applied Biosystems, Waltham, MA, United States) and TaqMan Gene Expression Assay (4331182 and 4351372, Applied Biosystems) (**Supplementary Table 14**) by the 7900HT System (Applied Biosystems). The reaction conditions were 95 °C for 10 min followed by 40 cycles of 95 °C for 15 s-60 °C for 1 min. 2 ^-ΔΔCt^ values^35^ were calculated from the obtained Ct values of the target and *Gapdh* mRNAs. For hypothalamus, *c-fos* mRNA was targeted, while in cultured cells the focus was on 16 mRNAs (**Supplementary Table 14**) related to the PI3K/Akt/mTOR pathway.

Reverse transcription and PCR of BDEV-derived miRNAs were also carried out according to the following procedure. Reverse transcription reactions were conducted using Custom RT primer pools mixed with 5× RT primer (4427975, TaqMan microRNA Assays, Applied Biosystems) and TaqMan microRNA reverse transcription kit (4366597, Applied Biosystems), with 150 pg of total RNA. The reaction conditions were 16 °C for 30 min, 42 °C for 30 min, and 85 °C for 5 min. Pre-amplification reactions were performed on the cDNA using TaqMan PreAmp Master Mix (4391128, Applied Biosystems). The PreAmp reaction conditions were 95 °C for 10 min, 55 °C for 2 min, 72 °C for 2 min, followed by 12 cycles of 95 °C for 15 s-60 °C for 4 min, and then 99.9 °C for 10 min. The reaction product was diluted 8-fold with 0.1×TE Buffer pH 8.0 (12090-015, Invitrogen). The PCR reactions were conducted using TaqMan Universal Master Mix II (4440040, Applied Biosystems) and 20× TaqMan microRNA Assays (4427975 and 4440886, TaqMan microRNA Assays, Applied Biosystems) (**Supplementary Table 15**) by the 7900HT

System (Applied Biosystems). The reaction conditions were 95 °C for 10 min, followed by 40 cycles of 95 °C for 15 s-60 °C for 1 min. 2^-ΔΔCt^ values were calculated using the Ct values of the target and internal standard (miR-16-5p^36^). The target miRNAs were 26 miRNAs (**Supplementary Table 15**) that changed more than 8-fold by microarray, had the same direction of change in the two pooled samples, and had human homologues.

### Behavioural test

To evaluate anxious behaviour, BDEVs or miRNA mimics were administered to mice immediately after acute stress exposure, and 24 h later, the open field test or the elevated plus maze test were performed on mice. Mice were acclimated to the testing room for at least 30 min prior to the start of the behavioural tests. All behavioural tests were conducted during 8:00-12:00.

Open field tests were conducted using a square (43.2 cm per side) open field (Med Associates Inc., Fairfax, VT, United States). Mice were placed in the centre, allowed to move freely for 15 min, and their activity levels at 10-15 min were analysed. The analysed behaviours were total distance travelled, a parameter of spontaneous locomotion, and centre time, an index of anxious behaviour. Analyses were performed using Activity Monitor Ver. 7 (Med Associates Inc.).

The elevated plus maze tests, using a maze with open and closed arms (650 mm long, 75 mm wide) placed 400 mm above the ground, were performed in a room illuminated at 20 lx. Mice were placed at the crossing, with their heads oriented towards the open-arm passage, and were allowed to move freely for 10 min. The ratio of the activity level in the open-arm area to the total activity level in all areas, a marker of anxiety behaviour, was analysed at 0-10 min. Analyses were conducted using AN Y-maze Ver. 7 (Stoelting Co., Wood Dale, IL, United States).

### *In vivo* administration

Intravenous administration of BDEVs or miRNA mimics was performed via the tail vein in unanesthetized mice placed in a fixture (28 mm diameter, 80 mm long) (ICN-2, ICM Corporation, Tsukuba, Japan) immediately after acute stress exposure. The dose of BDEVs purified from four mouse serum was 6 µg/0.1 mL Tris-glycine buffer/mouse. The preparation of the miRNA mimics was carried out according to the following procedure. A total of 60 µg of three custom miRNA mimics (20 µg each of miR-199a-3p, miR-99b-3p and miR-140-5p) (Horizon, Cambridge, United Kingdom) (**Supplementary Table 16**) were incubated with *in vivo* jet PEI (101000030; polyplus, Illkirch, France) and a 10% glucose solution (15 min, room temperature), and then formed miRNA-PEI complex was administered (60 µg/200 µL/mouse). The control mimic was cel-miR-67 (Horizon), a sequence from *C. elegans*.

Intracerebroventricular administration was conducted after a prior perforation and a subsequent recovery period. One week prior to administration, mice were underwent surgical removal of a small section of the skull and dura using a stereotaxic instrument. Post-surgery, a Dura-Gel (Cambridge NeuroTech, Cambridge, United Kingdom) was applied. One week after surgery, immediately after acute stress, BDEVs (0.15 µg/3 µL/mouse) purified from 24 mice serum and concentrated by ultracentrifugation was administered. The administration was carried out using syringe pump (Narishige, Tokyo, Japan) at a rate of 1 µL/min into the right lateral ventricle (bregma-0.4 mm, lateral 1 mm, depth 2.2 mm) of mice anaesthetized with isoflurane on a stereotaxic instrument.

### Cell culture

Mouse Neuro-2A (CCL-131) and mouse melanoma B16-F10 (CRL-6475) were purchased from ATCC (Manassas, VA, United States). Mouse microglia BV-2 was provided by Dr. Masahiro Ohgidani of Asahikawa Medical University.

Astrocyte isolation was performed according to a previous report^37^. Initially, the cerebral cortex of a P1 mouse was sampled and the meninges were removed under a stereomicroscope, while immersed in medium. Then, Stepwise tissue disruption was performed by suction ejection with needles of different gauges (18 G, 21 G and 23 G). After centrifugation (100 g, 5 min, room temperature), the supernatant was discarded. The obtained pellets were resuspended in medium and seeded into T-75 flasks coated with 0.5 mg/mL poly-D-lysine (150175, MP Biomedicals, Santa Ana, CA, United states). Once the astrocyte-containing cells had proliferated, contaminant cells were detached and removed by shaking the T-75 flask (200 rpm, 4 h, 37 °C). The adherent cells (astrocyte-rich) were detached using scrapers, then counted and seeded into T-25 flasks. In this astrocyte isolation method, neurons and oligodendrocytes are removed early in the purification process, while microglia may remain until late in the process. Hence, to confirm that no microglia remained in the isolated astrocytes, the expression level of the microglial marker *Iba1* mRNA was analysed. Consequently, *Iba1* expression in isolated astrocytes was 1/292.7 compared to BV-2, indicating high purity of astrocyte (**Extended Data Fig. 10**).

The culture mediums were as follows: for Neuro-2A, Minimum essential medium (11095098, Thermo), 1× Non-Essential Amino Acids (11140050, Thermo), 1 mM sodium pyruvate (11360070, Thermo), 10% deactivated fatal bovine serum (FBS) (SH30910.03, HyClone, Tokyo, Japan) and 100 U/mL Penicillin-Streptomycin (PS) (15140122, Thermo); for BV-2, Dulbecco’s modified eagle medium (DMEM) (high glucose) (11965092, Thermo), 10% deactivated FBS and 100 U/mL PS; for primary cultured astrocytes, DMEM/F12 (11320082, Thermo), 15% deactivated FBS and 100 U/mL PS. During astrocyte isolation, brain tissues were suspended in DMEM/F12 supplemented with 10% non-activated FBS and 1× Antibiotic-Antimycotic (15240062, Thermo). All cells were cultured at 37°C, 5% CO_2_.

### BDEV uptake to cultured cells

To demonstrate the uptake of BDEVs by neural cells, the following experiments were performed using the ExoSparkler Exosome Membrane Labeling Kit (Deep Red) (EX03, Dojindo, Mashiki, Japan) according to the instructions.

A mixture of Dye/DMSO solution with purified BDEVs (2-5 µg/100 µL) was incubated (37°C, 30 min). The labelled BDEVs solution was added to the column provided in the kit and centrifuged (3,000 g, 5 min, room temperature). Washing of the labelled BDEVs was then performed by addition of PBS and centrifugation (3,000 g, 5 min, room temperature). Finally, after the addition of PBS, the labeled BDEVs solution was collected. The negative control was a solution collected by conducting the same procedure using PBS.

Cells were seeded in 8-well coverglass chambers (5232-008, IWAKI, Yoshida-cho, Japan) 24 h prior to addition of labelled BDEVs. After PBS wash, 200 µL of medium and 50 µL of labelled BDEVs were added to the cells and incubated (37°C, 5% CO _2_, 4 h). Following incubation and a subsequent PBS wash, cells were examined under a fluorescence microscope (BZ-X710, Keyence, Osaka, Japan). Excitation and fluorescence wavelengths were 620/60 nm and 700/75 nm, respectively.

The medium was conditioned using exosome-free FBS (Exo-FBS-250A-1, System Bioscience, Palo Alto, CA, United States) and was PS-free. Cell seeding densities were set as follows: 1.0×10^6^ cells/mL for Neuro-2A and primary cultured astrocytes, and 1.0×10^5^ cells/mL for BV-2.

### Transfection to cultured cells (Transcriptome analysis)

Transient transfection of miRNA mimics into Neuro-2A, primary astrocytes and BV-2 was performed. To identify the target genes of the three *in vivo* administered miRNAs, a comprehensive analysis of mRNA expression was conducted.

The transfection medium was prepared as follows. Equal amounts of three custom miRNA mimics (miR-199a-3p, miR-99b-3p and miR-140-5p) (Horizon) (**Supplementary Table 16**) and Dharmafect 1 (T-2001, Horizon) were mixed to form a complex using PS-free serum-free medium. The mimic-Dharmafect1 complex were then diluted with complete medium to 25 nM. For the control mimic, cel-miR-67 was used (Horizon).

After cultivation in PS-free medium (37°C, 5% CO_2_, 24 h), cells were incubated in transfection medium for 24 h. Following transfection, RNA was extracted using QIAzol lysis reagent. Finally, obtained RNA samples were analysed by NGS (DNA Chip Research Institute, Tokyo, Japan). Cell seeding was conducted at the following concentrations: Neuro-2A at 6.0×10^5^ cells/mL, primary astrocyte at 9.0×10^5^ cells/mL and BV-2 at 3.0×10^5^ cells/mL.

### *In silico* analysis

Candidate mRNAs targeted by miRNAs were selected from the down-regulated genes identified by transcriptome analysis in the following manner. Since it was not known whether the changes in the down-regulated genes by the three miRNAs (miR-199a-3p, miR-99b-3p and miR-140-5p) were direct or indirect, the down-regulated genes were matched to the miRNA-targeted mRNAs identified using TargetScanMouse Ver. 8.0 (https://www.targetscan.org/mmu_80/). As a result, genes extracted by both the down-regulated genes and TargetScanMouse were selected as direct target candidates. In addition, Upstream Regulator analysis by Ingenuity Pathway Analysis (Qiagen) was performed to determine the contribution of each of the three miRNAs (miR-199a-3p, miR-99b-3p and miR-140-5p) to the down-regulated genes. For identifying behavioural factors, “behaviour”-related categories were extracted using Diseases or Functions Annotation analysis of Ingenuity Pathway Analysis. Then factors within each category were elicited.

### 3′ UTR reporter assay

The 3’ UTR of *Mecp2*, containing the target sequence (positions 285 to 292) as predicted by TargetScanMouse, was cloned from total RNA of Neuro-2A using PCR. The following primers were used for cloning: Forward: 5’-CACTCGAGTTCTTTAC ATAGAGCGG ATTGC-3’ including Xho I restriction site, Reverse: 5’-CAGTCGACTACCTTGTCAGTTTTGG ATTGG-3’ including Sal I restriction site. 3’polyadenylate overhang was added to the PCR products after treatment with Taq polymerase (72°C, 15 min). The cloned into a pGEM-T easy vector (A137A, Promega, Madison, WI, United States). The amplified products were ligated into the Xho I and Sal I sites of the 3’UTR of the firefly luciferase gene in the pmirGLO Dual-Luciferase miRNA Target Expression vector (E 1330, Promega) to generate pmiR-Mecp2. Site-directed mutagenesis was performed in the seed sequences of Mecp2 using PrimeStar Max DNA Polymerase (R045, Takara, Japan) for PCR amplification. The following primers were used for site-directed mutagenesis: Forward: 5’-AG ATCCGA ATTACCAGACAAGCTGTTGACC-3’ Reverse: 5’-TGGTA ATTCGG ATCTGCAGCAAGCCTTGT-3’. HEK293 cells were simultaneously transfected with either wild type or mutant pmi R-Mecp2 vectors containing the 3’UTR, along with miR-199a-3p mimic or negative control mimic. Subsequently, firefly and renilla luciferase activities were measured. Transfection and assay conditions are shown below. Cells were seeded in 96-well plates at a concentration of 1.0 × 10^4^ cells/well and incubated for 24 h. Then, pmi R-Mecp2 vector or pmiR-Mecp2 with mutation in seed sequence vector and mimic were transfected into the cells using DharmaFECT Duo reagent (T-2010, Horizon). At 48 h after transfection, cells were harvested and firefly luciferase activity values were normalized using renilla luciferase activity values.

### Statistical analysis

Statistical analysis was performed using JMP Pro 15.2.0 (SAS Institute Inc., Cary, NC, United States). For comparisons between two groups, an independent t-test was used, while ANO VA followed by Tukey-Kramer post hoc test was utilised for comparisons among three or more groups. Both tests were two-tailed and set at a significance level of 0.05. The Benjamini-hochberg method was used for NGS analysis results.

---

**Extended Data Fig. 1.**
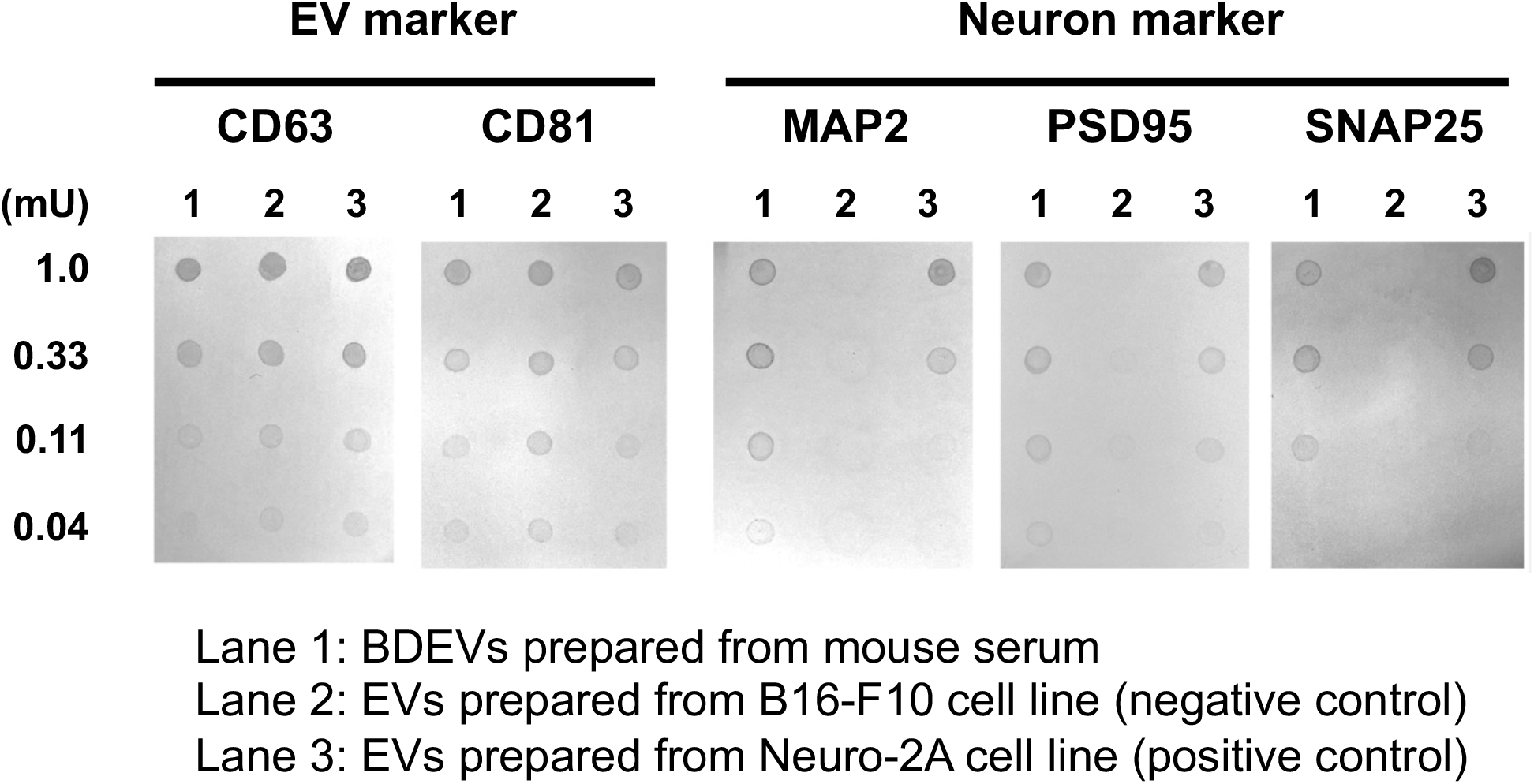
BDEVs isolated from serum is of neuronal origin. Signals of EV and neuronal markers in BDEV isolated from serum were observed, and they exhibited a dose-dependent manner.

**Extended Data Fig. 2.**
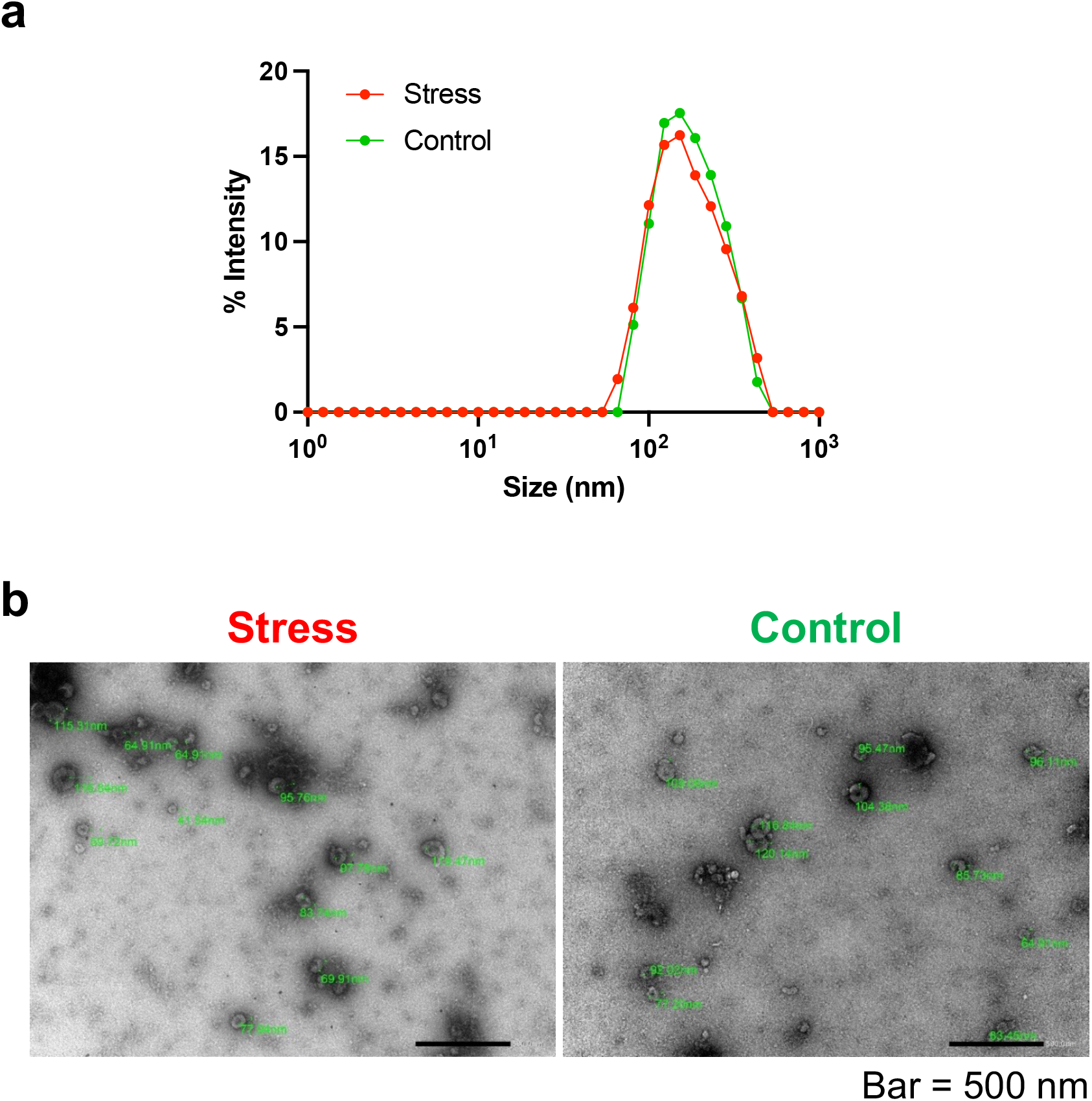
Acute stress does not affect EV appearance. **a**, Acute stress did not alter EV size distribution. **b**, Acute stress did not alter EV morphology. EV, extracellular vesicles.

**Extended Data Fig. 3.**
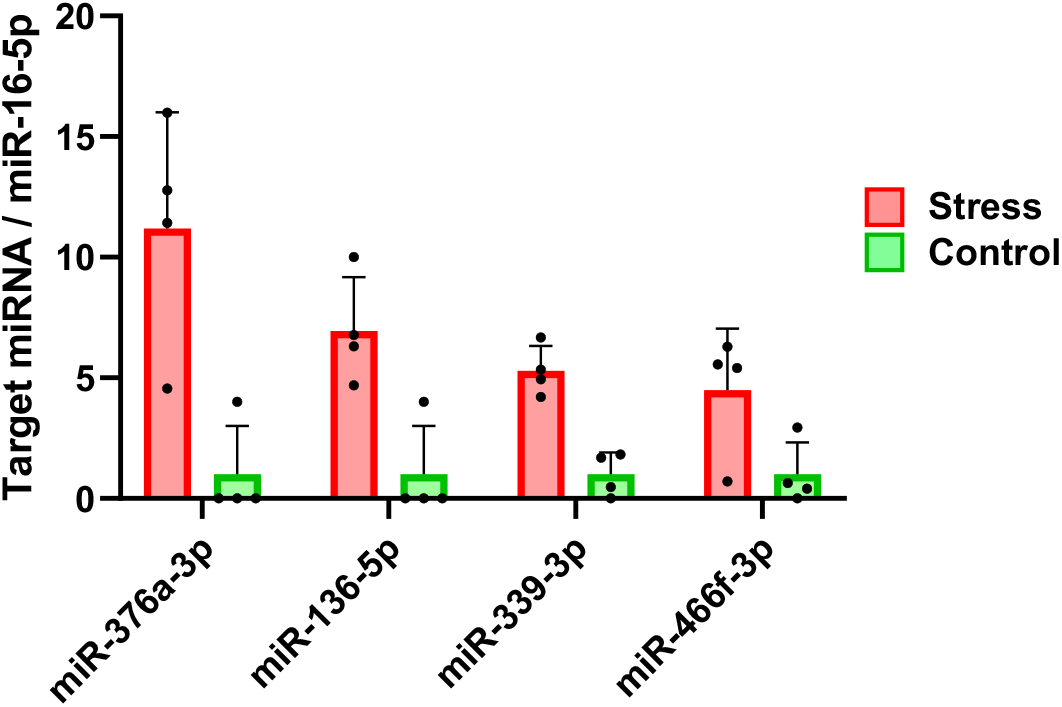
About the four BDEV-encapsulated miRNAs excluded from further analysis. Four miRNAs shown here (miR-376a-3p, miR-136-5p, miR-339-3p and miR-466f-3p) were included in the 26 miRNAs shown in Fig. 2f. However, real-time RT-PCR showed that the Ct values of any of the miRNAs in the control group was below the measurable sensitivity, so the relative expression levels of the miRNAs could not be determined (n=4 pools, each consisting of 2 mice). Data are mean±SD. BDEV, Brain-derived extracellular vesicles.

**Extended Data Fig. 4.**
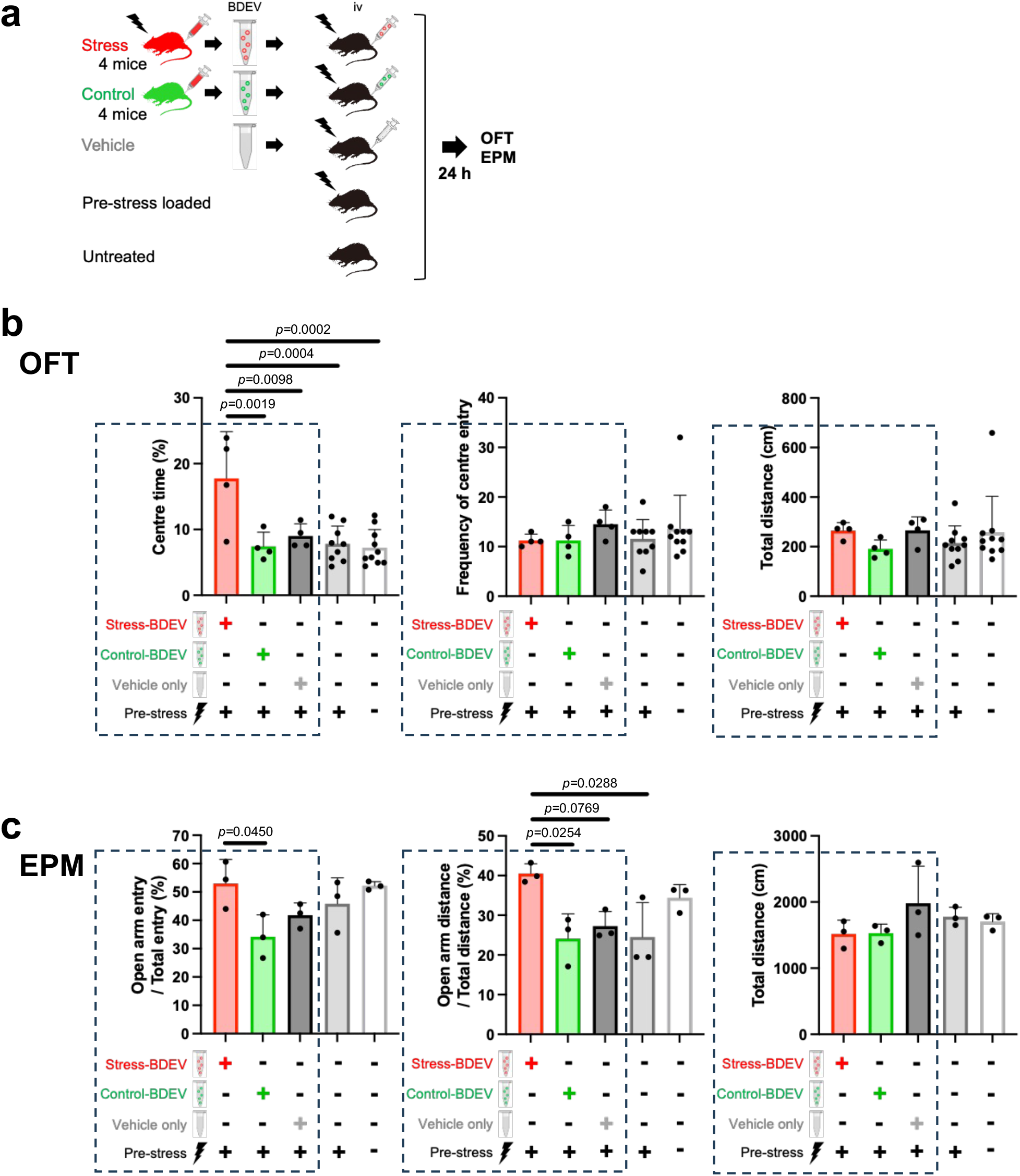
Stress-induced BDEVs ameliorate anxiety behaviour. **a**, The behavioural tests (OFT and EPM) shown in **Fig. 3a,c** were actually conducted with five groups. The BDEVs were administered 24 h after the acute stress exposure (pre-stress loaded) and the behavioural tests were conducted immediately thereafter. Therefore, to ensure that there was no effect of pre-stress on the behavioural tests, a pre-stress only group without intravenous injection (pre-stress loaded group) and a pre-stress unloaded group without intravenous injection (untreated group) were added and their behavioural levels were assessed. **b**, Evaluation by the open field test. Intravenous administration of stress-BDEVs (red) significantly increased the centre time at 10-15 min after the start of the open field test compared to the other four groups (n=4-10). In contrast, the frequency of centre entry and the total distance traveled did not differ between any of the groups. A portion of each graph was extracted (within the *dashed line*) and displayed as **Fig. 3b. c**, Evaluation by the elevated plus maze test. Intravenous administration of stress BDEVs (red) significantly increased the frequency of open arm entry compared to the control-BDEVs (green) (n=3) and significantly increased the distance travelled in the open arm compared to the control-BDEVs (green), vehicle (dark grey) and stress (light grey) groups (n=3). Conversely, no difference in total distance was found between any of the groups. A portion of each graph was extracted (within the *dashed line*) and displayed as **Fig. 3d**. Data are mean±SD. Statistical analysis was performed using one-way ANOVA with Tukey-Kramer post hoc test. BDEV, Brain-derived extracellular vesicles.

**Extended Data Fig. 5.**
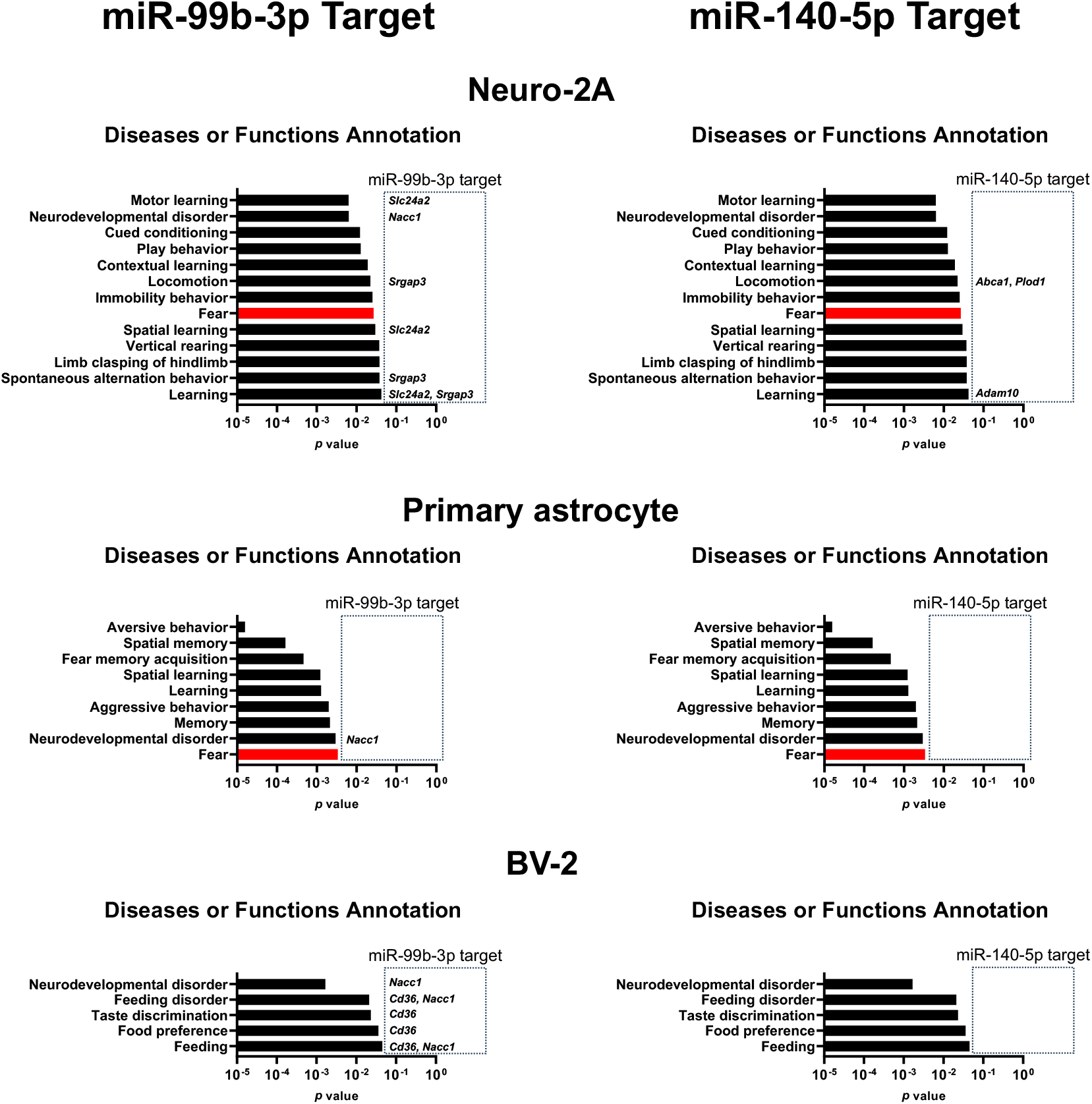
No target genes for miR-99b-3p and miR-140-5p are included in the “fear” category in the IPA. Categories of significantly changed behaviour as identified by IPA’s Disease or Function Annotation analysis. And the mRNAs shown within the *dashed lines* are those associated with each category and identified as targets of miR-99b-3p or miR-140-5p by TargetScanMouse (Supplementary Table 5,6). No genes targeted by miR-99b-3p or miR-140-5p appeared in the ‘Fear’ category. Statistical analysis was performed using the binomial test. IPA, Ingenuity pathway analysis.

**Extended Data Fig. 6.**
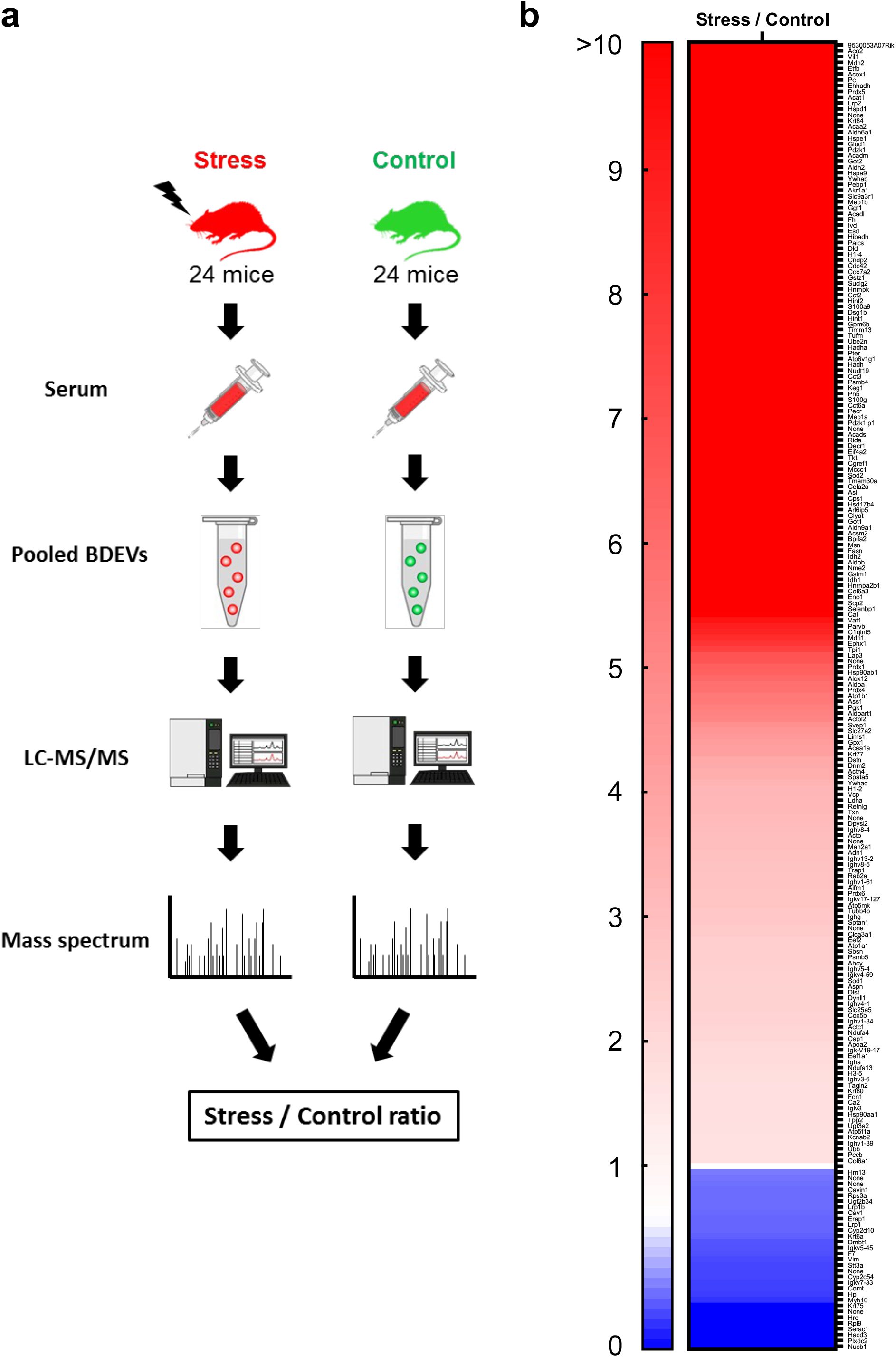
Acute stress alters protein profile of BDEVs. **a**, Schematic protocol of proteome analysis in BDEVs using LC-MS/MS. **b**, Proteins up- and down-regulated by acute stress. Of the 803 proteins identified by proteome analysis, 193 proteins showed an increase of more than 2 (shown in *red*) and 31 proteins decreased to less than 0.5 (shown in *blue*). (See supplementary table 9 for full details).

**Extended Data Fig. 7.**
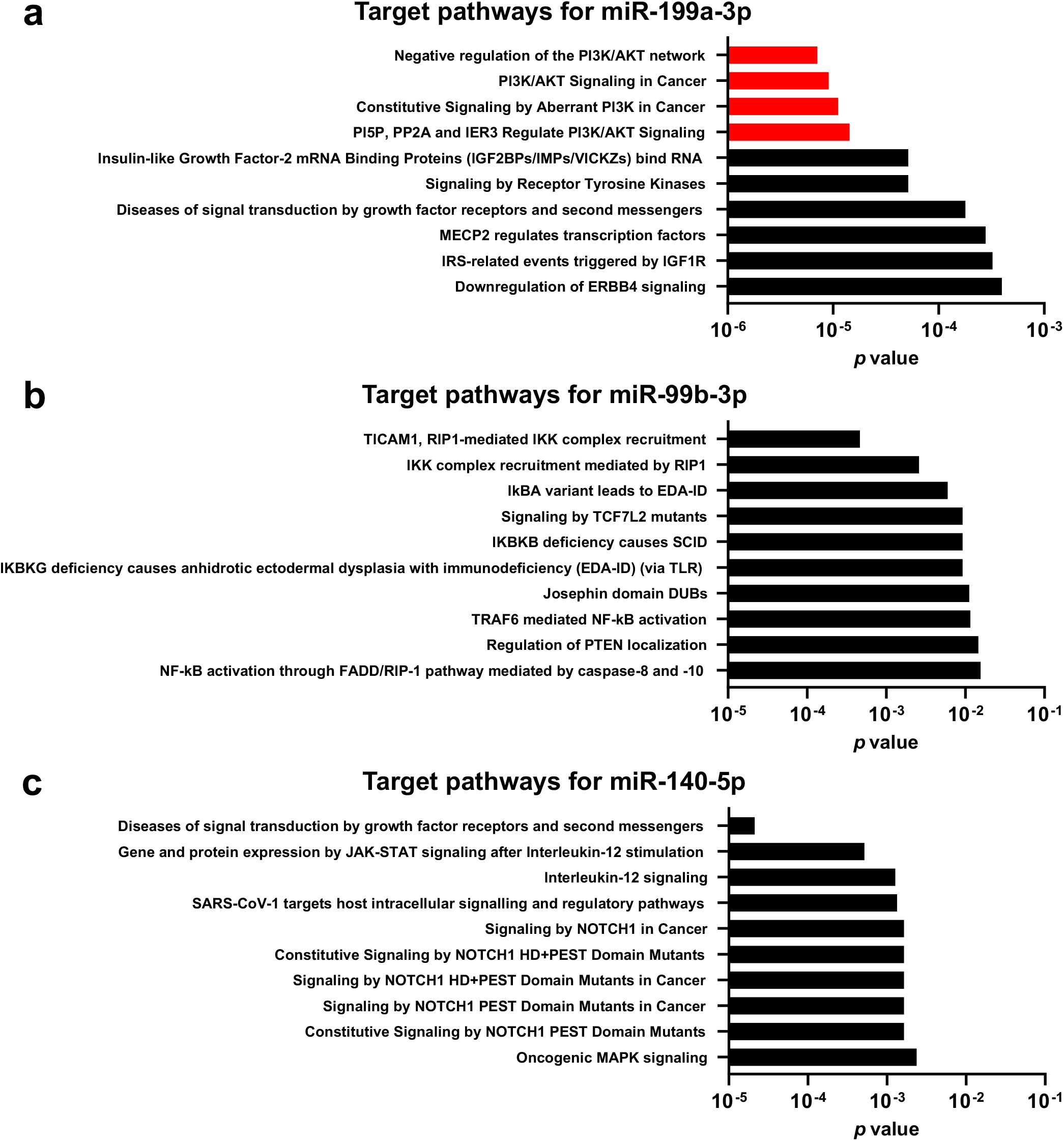
Top 10 target pathways of stress-responsive BDEV-encapsulated miRNAs by *in silico* analysis. **a, b, c**, Target pathways for the three miRNAs. The three miRNA target candidate genes identified by TargetScanMouse were categorised using the Reactome Pathway Database. For miR-199a-3p (**a**), the top 4 cascades (coloured red) were all PI3K/Akt/mTOR pathways, however, for miR-99b-3p (**b**) and miR-140-5p (**c**), PI3K/Akt/mTOR-related pathways were not included. Statistical analysis was performed using binomial test.

**Extended Data Fig. 8.**
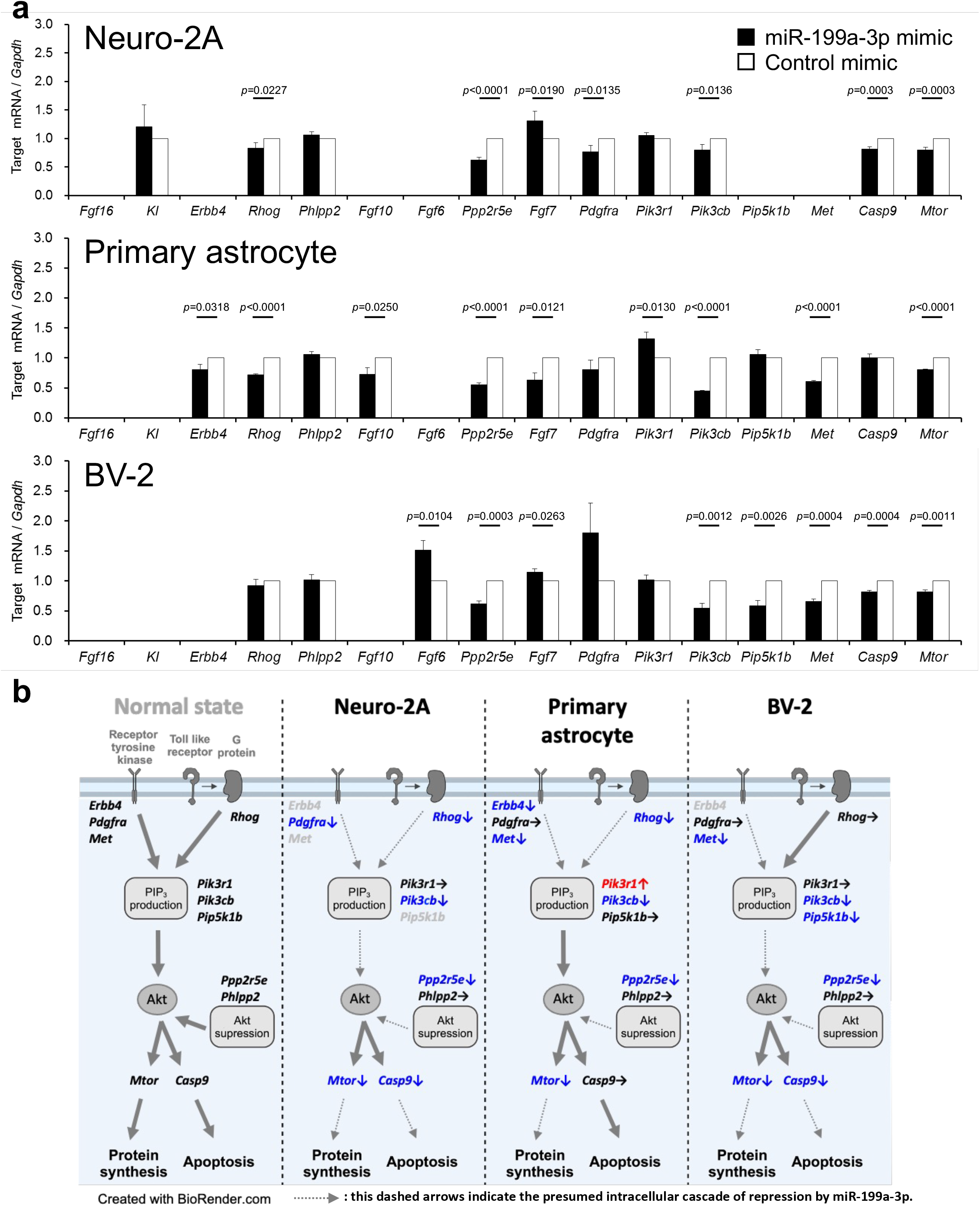
Stress induced miR-199a-3p suppresses PI3K/Akt/mTOR pathway in neural cells. **a**, Changes in mRNA belonging to the PI3K/Akt/mTOR pathway when miR-199a-3p is added to each cell. More than half of the PI3K/Akt/mTOR pathway-related genes are significantly altered in all three cells. (Neuro-2A; n=4, Primary astrocyte; n=3, BV-2; n=3). **b**, Mapping of factors shown in “***a***” on the pathway. Italics indicate factors analysed in this study. Grey coloured genes were not detected. Dashed grey arrows indicate pathways with presumed reduced activity. Coloured arrows are: blue; decreased expression, red; increased expression, black; no change. Data are mean±SD. Statistical analysis was performed using Student’s two-tailed *t*-tests.

**Extended Data Fig. 9.**
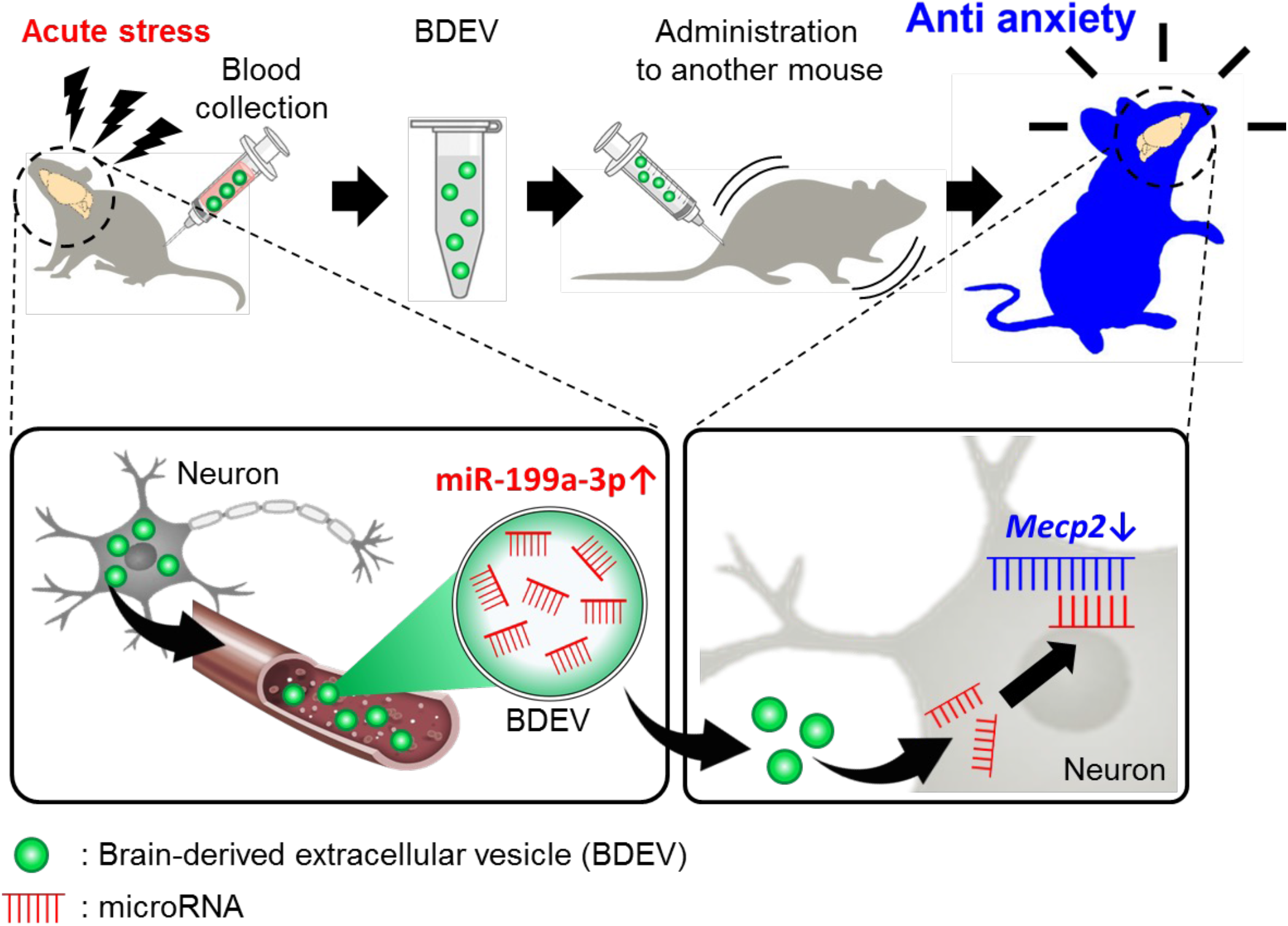
Graphical summary of this study.

**Extended Data Fig. 10.**
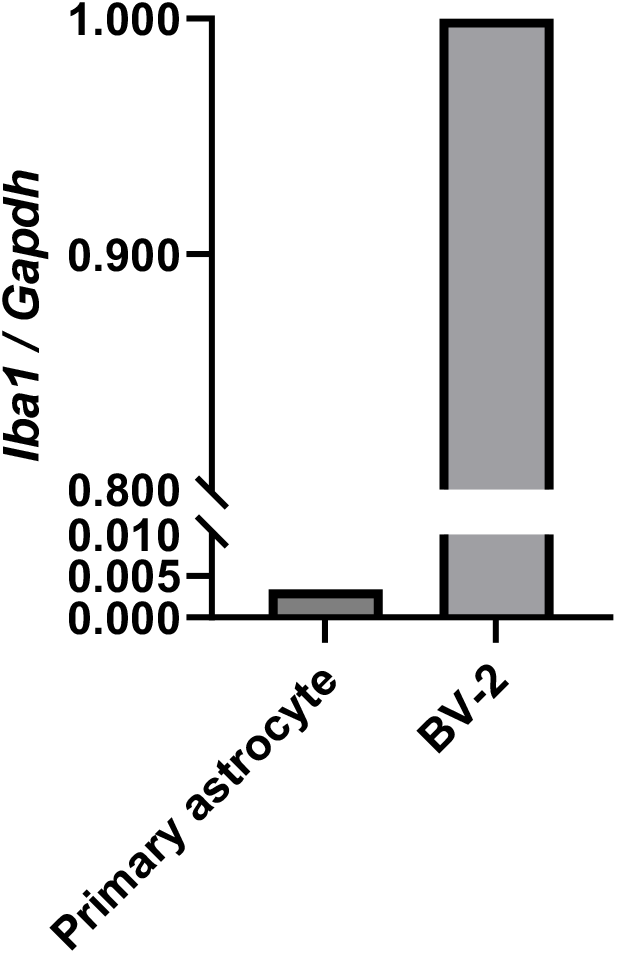
Isolated primary cultured astrocytes contain almost no microglial cells. The expression of the microglial marker *Iba1* in isolated astrocytes was compared with that in mouse microglial cell line BV-2 by qPCR. The results showed that the expression of *Iba1* in astrocytes was 1/270 of that in BV-2, indicating that astrocytes were highly purified.

## Notes

### Competing Interest Statement

The authors have declared no competing interest.

